# Benchmarking AI-based plasmid annotation tools for antibiotic resistance genes mining from metagenome of the Virilla River, Costa Rica

**DOI:** 10.1101/2023.08.24.554652

**Authors:** Dorian Rojas-Villalta, Melany Calderón-Osorno, Kenia Barrantes-Jiménez, Maria Arias-Andres, Keilor Rojas-Jiménez

## Abstract

Bioinformatics and Artificial Intelligence (AI) stand as rapidly evolving tools that have facilitated the annotation of mobile genetic elements (MGEs), enabling the prediction of health risk factors in polluted environments, such as antibiotic resistance genes (ARGs). This study aims to assess the performance of four AI-based plasmid annotation tools (Plasflow, Platon, RFPlasmid, and PlasForest) by employing defined performance parameters for the identification of ARGs in the metagenome of one sediment sample obtained from the Virilla River, Costa Rica. We extracted and sequenced complete DNA from the sample, assembled the metagenome, and then performed the plasmid prediction with each bioinformatic tool, and the ARGs annotation using the Resistance Gene Identifier web portal. Sensitivity, specificity, precision, negative predictive value, accuracy, and F1score were calculated for each ARGs prediction result of the evaluated plasmidomes. Notably, Platon emerged as the highest performer among the assessed tools, exhibiting exceptional scores. Conversely, Plasflow seems to face difficulties distinguishing between chromosomal and plasmid sequences, while PlasForest has encountered limitations when handling small contigs. RFPlasmid displayed diminished specificity and was outperformed by its taxon-dependent workflow. We recommend the adoption of Platon as the preferred bioinformatic tool for resistome investigations in the taxon-independent environmental metagenomic domain. Meanwhile, RFPlasmid presents a compelling choice for taxon-dependent prediction due to its exclusive incorporation of this approach. We expect that the results of this study serve as a guiding resource in selecting AI-based tools for accurately predicting the plasmidome and its associated genes.

## I. Introduction

Rivers are relevant water sources for human development, providing recreation, tourism, agriculture, and electricity, among other resources. However, anthropogenic activities regarding sources exploitation and inadequate regulation and treatment of wastewater have increased the contamination of water bodies [1]. In Costa Rica, despite their low efficiency, septic tanks remain the primary sanitary method for wastewater treatment; with less than a quarter connected to sewerage, and only 15.5% of the total incoming is treated before being discharged into rivers [2].

Previous studies have shown aggravated contamination near areas with accelerated socio-economic development, due to the lack of efficient wastewater treatment [3]. In this sense, the Great Metropolitan Area of Costa Rica (GAM) represents a potential source of water contamination as it contains the most significant industrial and commercial sector in the country [4]. The Virilla River Basin drains this area, receiving approximately 67% of the country’s wastewater and being considered the most polluted river in Central America [4]. Among many pollutants, multi-residue analysis of samples has shown the presence of pharmaceutical compounds in the Virilla River (e.g. antibiotics: ofloxacin, cephalexin) [5]. These compounds represent a clinical risk as they might promote the dissemination of antibiotic resistance genes among bacteria, including pathogenic species [6], [7].

At the same rate as the different -omics sciences have progressed, the study of the occurrence, abundance, and diversity of ARGs has been facilitated [8], [9]. This -omics approach is extensively used to evaluate ARGs in aquatic environments, showing an advantage in comparison with culture-dependent techniques by identifying the resistome (all ARGs in a microbial community) composition of non-culturable microorganisms [9]–[13]. Many bioinformatic tools are currently established for the annotation of mobile genetics elements, specifically the plasmidome (all plasmids in a microbial community) as they represent the main vector for horizontal gene transfer (HGT) of ARGs and other traits [7]. In addition, the implementation of artificial intelligence (AI) and machine learning algorithms (such as random forest and artificial neural networks) have successfully improved metagenomic data analysis favorably in its precision and speed, including the annotation of MGEs [14].

Here, we provide a benchmarking analysis between four different AI-based bioinformatic tools (Plasflow, Platon, RFPlasmid, and PlasForest) for plasmids annotation, based on the performance for the identification of antibiotic resistance genes in a sediment metagenome from the Virilla River (San José, Costa Rica).

## II. Materials and Methods

### A. Sample collection, DNA extraction, and sequencing

Sediment and surface water samples were collected from Virilla River (San José, Costa Rica) on November 1st, 2021 (Coronado city, 9°59’09.1”N 83°56’35.0”W, 2020 meters ASML). For sediments, a sterile spoon was employed to collect samples from the river’s shore under running water at a depth of approximately 30 cm. The collected sample was carefully placed into a sterile bag and subsequently sealed within another sterile bag to maintain its integrity. Four liters of surface water were collected using a sterile recipient. Both types of samples were maintained in a cold environment during transportation to the laboratory. Dissolved organic carbon (DOC), fecal coliforms, and oxygen saturation were analyzed to determine the surface water quality based on [15], and a water quality index as mentioned by [16]. All assays were done in triplicate and obtained values were averaged. For sediment samples, complete DNA was extracted by Power Soil Pro Extraction Kit (QIAGEN) and then sequenced through a shotgun metagenomics approach using an Illumina NovaSeq PE150 platform (Novogene; California, USA).

### B. Metagenome assembly and data management

The quality assessment of raw Illumina reads was conducted using FastQC v0.11.7 [17]. To eliminate low-quality and unpaired reads, we applied Trimmomatic v0.36, [18] considering Phred-33 quality scores with an average value of 35. Subsequently, the filtered and paired raw reads were subjected to assembly using metaSPAdes v3.12.1 [19] with default parameters, employing K-mer values of 55, 65, and 75, and utilizing 72 threads for enhanced processing efficiency. Following the plasmid annotation software’s suggestion, contigs with a length of less than 1000 bp were excluded using the “reformat.sh” script from bbmap v37.36 [20]. The quality assessment of the filtered assembly was performed using QUAST v4.6.0. [21].

### C. Plasmid sequences annotation by AI-based tools

We selected the four bioinformatic programs leveraging AI and machine learning released (to our knowledge) at the time this research was being conducted: PlasFlow v1.1 [22], Platon v1.6 [23], RFPlasmid v0.0.18 [24], and PlasForest v1.2 [25].

Overall, PlasFlow depured a dataset of chromosomal/plasmid sequences and transformed it as Term Frequency-Inverse Document Frequency (TF-IDF; Scikit-learn v0.18 [26]) vectors from frequencies of all oligonucleotides (k-mer) of desired length (k): genomic signature k-mers (k value: 3 to 7) [22]. Normalized vectors were used to train the neural network with TensorFlow^1^ until its final model after 50,000 learning steps [22].

Platon calculated replicon distribution scores (RDS) from a database of marker protein sequences from bacterial sequences based on frequencies of being encoded in plasmids or chromosomes [23]. A higher-level characterization using plasmid-related sequence analysis was combined with RDS for the final learning Monte Carlo algorithm of the tool, trained with the initial database [23].

RFPlasmid utilized genomic signature k-mers (k value: 5) from all plasmid and chromosome sequences reported for 17 bacterial genera individually; the agnostic approach (the one used in this study) database was generated by concatenating all the datasets [24]. Random Forest [26] models for 17 individual genera and agnostic approach were trained using 5,000 trees [24].

Finally, PlasForest employed a homology-based methodology. Bacterial genomes artificially recreated as the empirical distribution of contigs sizes of unassembled genomes were compared using BLASTn against a downloaded plasmid sequence database [25]. Extracted genomic features from “hits” and contigs dataset were used to train the Random Forest Classifier algorithm [26] with 500 trees [25].

It is relevant to remark that these tools were developed using different algorithms, databases, and methodologies. However, published final models of the programs don’t use input files for learning, instead, these are utilized for common annotation pipelines in the already-learned tool. Therefore, no hyperparameters of the AI algorithms are modified in this study. We aimed purely to benchmark the tools’ annotation performance on environmental metagenomic data based on posterior analyses, from a biological perspective.

For this, the programs were executed on the same computational allocation, utilizing nodes equipped with Intel Xeon processors having 36 cores with 2 threads each @ 3.00 GHz and 1024 GB of RAM and parallelization across the available 72 threads. All bioinformatic tools were run using their default parameters as there are no common flags among the programs and output prediction wouldn’t be comparable if used. For RFPlasmid, we employed the web portal version^2^, and the ‘Generic’ species already-learned database was specified. Subsequently, the annotated plasmidomes’ quality was evaluated using QUAST v4.6.0 [21].

### D. Functional annotation for antibiotic resistance genes

Plasmid sequences were annotated using the Resistance Gene Identifier v6.0.1 web portal from the Comprehensive Antibiotic Resistance Database v3.2.6 (CARD)^3^ [27]. The process involved Open Reading Frame prediction with Prodigal, homologs detection using DIAMOND, and strict significance annotation based on the curated cut-off bitscores of CARD [27]. Additionally, we used the whole metagenome as a positive control for benchmarking analysis. To ensure inclusivity, we set the sequence quality and coverage to low, thereby avoiding the exclusion of small plasmids (less than 20,000 bp) and some assembled contigs.

### E. Performance analysis

We categorized antibiotic resistance genes (ARGs) found in plasmidomes as the positive class, while ARGs exclusively predicted in the whole metagenome were considered the negative class. Each AI-based tool’s prediction for the annotated plasmidome was compared against the resistome of the entire metagenome. Resistance gene prediction results were assigned to the following categories: i) True Positive (TP), plasmid-associated ARG predicted in both the evaluated plasmidome and the whole metagenome, ii) True Negative (TN), chromosome-associated ARG predicted only in the whole metagenome, iii) False Positive (FP), chromosome-associated ARG predicted in the plasmidome, and iv) False Negative (FN), plasmid-associated ARG not predicted in the evaluated plasmidome. We then calculated the performance parameters of Sensitivity, Specificity, Precision, Negative Predictive Value, Accuracy, and F1-score, considering ARGs prediction as 1 when present and 0 when not found. ARGs association was determined according to a literature review of previously reported presence in MGEs and chromosomal-integration events, mentioned in the results and discussion’s second subsection.

## III. Results and discussion

### A. Assembled whole metagenome and annotated plasmidomes through AI-based tools

Chemical and biological analysis of the water sample revealed a 1.183 DOC value, 7.74 O_2_ mg/L, 288 MPN/100 mL for fecal coliforms, and NSFQI quality of 66 based on [16]. This indicates that pollution is moderate at this sampling site at the Virilla River. The assembled metagenome displayed a length of 106.5 Mbp, consisting of 45,471 contigs. For the plasmidomes, PlasFlow annotated 22.9 Mbp (11,883 contigs), Platon identified 0.2 Mbp (55 contigs), RFPlasmid revealed 0.5 Mbp (245 contigs), and PlasForest predicted 0.1 Mbp (88 contigs). Additional characteristics are provided in Table I.

**TABLE 1.** Genomic characteristics and quality parameters of assembled whole metagenome and annotated plasmidomes.

|  | Whole | Plasflow | Platon | RFPlasmid | PlasForest |
| --- | --- | --- | --- | --- | --- |
| Size | 106 | 22.9 | 0.2 | 0.5 | 0.1 |
| Contigs | 45471 | 11883 | 55 | 245 | 88 |
| N50 | 2592 | 1924 | 4374 | 2487 | 1475 |
| L50 | 9672 | 3218 | 11 | 51 | 31 |

Size is presented in Mbp

The number of plasmids found in metagenomes varies according to the sample source. In a swine manure treatment plant sample, the alignment against a database built of all plasmid sequences reported in the NCBI, demonstrated the presence of 59 to 440 plasmids [28]. Other authors evidenced the presence of 5611 and 7184 plasmids from two plasmidome water samples of Jeju Island (South Korea) [29]. As mentioned, the Virilla River samples are considered moderately polluted, therefore the number of plasmids might be similar to those found in other polluted environments, such as the treatment plant. Moreover, the size and number of plasmids annotated by Plasflow are widely different from those observed in other plasmidomes, which presume the idea of possible chromosome contigs incorrectly identified as plasmids by the bioinformatic tool, as previously demonstrated [23].

### B. Resistome annotation of whole metagenome and plasmidomes predicted by AI-based tools

The whole metagenome annotation revealed the presence of 21 antibiotic resistance genes (ARGs), including mainly resistance against disinfecting and antiseptics agents, fluoroquinolone, tetracycline, glycopeptide, sulfonamide, and aminoglycoside antibiotics (Supplementary Table I). Predicted plasmids showed divergence in their results regarding ARGs quantity. Plasflow-, Platon-, RFPlasmid-, and PlasForest-annotated plasmidomes presented six, four, three, and four of the total ARGs annotated, respectively (Supplementary Table I). Remarkably, three resistance genes had the highest similarities to the matching region and were present in all annotations: *qacEdelta1* (100%), *sul1* (99.64%), and *APH(3”)-lb* (99.25%).

We observed a remarkable number of ARGs regarding disinfecting and antiseptics agents, fluoroquinolone, and tetracycline antibiotics. In the past few years, quinolones resistance has been an increasing problem due to the high possibility of HGT demonstrated in conjugation studies [30], [31]. Chemical analysis of water samples from Virilla River showed the presence of residues of antibiotic compounds from the fluoroquinolone family (ofloxacin) [5], which might represent a selective pressure for HGT [32]. In addition, these ARGs are known to be associated with sewage water bacteria [31].

However, most of these ARGs are not annotated in the plasmidomes. Fluoroquinolone resistance genes are common in the genome of pathogenic bacteria [32]. Integron analysis revealed the association of these ARGs to genetic elements possibly acquired by plasmids or other chromosomes [33]. The resistance against aminoglycosides, macrolides, tetracycline, and beta-lactam was also observed with a similar pattern [33]. These results can indicate the possibility of our findings regarding similar resistance genes to be integrated into chromosomes of the metagenome, explaining their absence in the plasmidomes. This hypothesis is supported by other genomic studies where the ARGs related to fluoroquinolone and tetracycline resistance are associated with the bacterial chromosome [34], [35].

Moreover, annotated disinfecting and antiseptics agents, and glycopeptide antibiotics resistance genes (mainly found in the whole metagenome) are associated with a high rate of horizontal transfer [36]. These ARGs have been demonstrated to also be ubiquitous in the chromosomal genomes [37], elucidating the idea of their integration into the chromosomes of the metagenome. In addition, the presence of tetracycline resistance could be associated with biocide-reduced susceptibility [38], unifying the presence of both ARGs annotation into metagenomic chromosomes and absence in plasmidomes. Finally, the same pattern is reported for genes related to glycopeptide resistance, including those annotated in our metagenome but not in the plasmidomes [39].

Low concentrations of antibiotics in the environment can promote resistance due to antibiotic resistance genes [40]. As mentioned, traces of these pharmaceutical compounds such as ofloxacin and cephalexin have been found in the Virilla River [5], representing a sanitary risk for microbial communities, ecological relations, and major organism aquatic organisms [41]–[43]. ARGs are commonly mobilized between bacteria via plasmids in environment-related processes of HGT (e.g., conjugation and transformation) [7], [44]. The presence of pollutants in aquatic environments is another factor involved in the aggravation of ARGs’ dissemination [7], [45]. Therefore, studying the resistome present in the Virilla River is a relevant research topic to elucidate the significant risk of inefficient wastewater treatment methods and their effect on the worldwide clinical antibiotic resistance crisis.

### C. Performance evaluation of AI-based plasmid annotation tools in terms of ARGs prediction

As for the benchmarking analysis of the plasmid annotation tools, the presence of glycopeptide, fluoroquinolone, and tetracycline antibiotics resistance genes are considered False Positives due to the statements previously discussed. These ARGs are mainly found in the chromosome of microorganisms and were only observed in Plasflow- and PlasForest-annotated plasmidome, therefore we evaluated them as a precision and accuracy discrepancy between AI-based programs. The distribution of ARGs and score for each results category per plasmidome sample is shown in Fig.1.

**Fig. 1.**
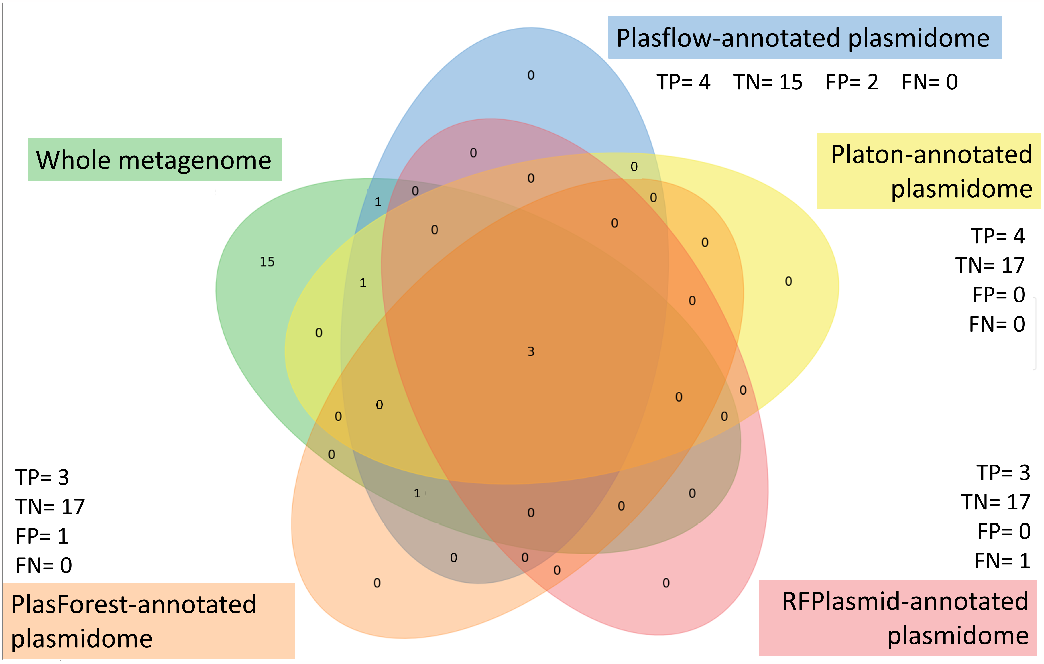
Venn’s diagrams of the number of predicted ARGs per sample and score for each results category per plasmidome annotated with AI-based tools. TP=true positive, TN=true negative, FP=false positive, FN=false negative.

Platon presented the highest scores regarding the performance metrics, followed by RFPlasmid and PlasForest (Table II). Plasflow showed the lowest precision and accuracy among the annotation tools, which correlates with previous studies that indicate an imprecise differentiation between bacterial chromosomal sequences from plasmids [23]. These results explain the False Positive ARGs found in the resistome, as well as the size and number of contigs differences of the Plasflow-annotated plasmidome in comparison with the other annotation tools, stated in the first subsection. Moreover, PlasForest has been reported to have less sensitivity and precision for plasmids identification when working with small contigs (less than 2000 bp), being surpassed even by Plasflow [25]. As the whole metagenome was filtered to 1000 bp as the minimum contigs length, the false positives outputted by PlasForest might be a result of its difficulty in annotating small sequences.

**TABLE 2.** Performance metrics for Plasflow, Platon, RFPlasmid, and PlasForest plasmidomes annotation based on prediction of antibiotic resistance genes.

| Metric | AI-based plasmid annotation tools |  |  |  |
| --- | --- | --- | --- | --- |
|  | Plasflow | Platon | RFPlasmid | PlasForest |
| Sensitivity (%) | **100** | **100** | 75 | **100** |
| Specificity (%) | 88.23 | **100** | **100** | 94.44 |
| Precision (%) | 66.67 | **100** | **100** | 75 |
| NPV (%) | **100** | **100** | 94.44 | **100** |
| Accuracy (%) | 90.47 | **100** | 95.23 | 95.23 |
| F1-score (%) | 80 | **100** | 85.71 | 85.71 |

Bold text represents the highest value. NPV=Negative Predictive Value

RFPlasmid metrics evidence the lowest sensitivity. Regardless of presenting an accuracy within the acceptable range, the “Generic” model (taxon-independent workflow) used for predicting has been reported to be outperformed by its taxon-dependent workflow in various studies [24], [46]. All the bioinformatic tools here evaluated used a taxon-independent approach, therefore, usage of RFPlasmid under this agnostic model for metagenomic exploratory research could incur incomplete plasmidomes annotation and inaccurate results [46]. While undergoing literature revisions, we found a lack of information regarding the benchmarking of AI-based annotation tools. However, previous research involving the performance evaluation of plasmids prediction tools, including those here inspected, showed similar performance metrics patterns. Despite presenting the lowest number of contigs, Platon had the highest scores (near 100%) in comparison with other examined tools, while Plasflow remained the least precise and accurate program for plasmidome annotation in environmental metagenomic datasets [46], [47].

Platon integrates, in its higher-level classification, the detection of antimicrobial resistance genes [23]. This might explain the high performance of this study as it incurs overfitting of the identification process of the tool and the data analyzed for performance evaluation. In addition, the Platon dataset for training is based on processed plasmid sequences, different from other programs [22]–[25]. The dataset selection, combined with plasmid-specific heuristics parameters, might confer Platon a more detailed detection capacity of plasmid sequences.

This eludes that Platon differentiates chromosomal and plasmid sequences with more accuracy, especially when ARGs are present, outputting a small, but more precise, plasmidome [23]. PlasForest presented similar metrics, although is outperformed by Platon when working with small contigs. We recommend Platon for resistome research with exploratory objectives as it presented the best performance in plasmids prediction, while RFPlasmid could represent a better suit for species-specific research due to its taxon-dependent workflow, unique among the evaluated tools.

Plasmids play a crucial role in bacterial genetics, facilitating the mobilization of genetic elements for microorganisms’ adaptation to the environment, including antibiotic resistance genes. We explored the presence of ARGs as well as their mobilization by HGT. Despite their relevance and the variety of methods for plasmid annotation, there is no standard method to compare the accuracy; leading to a lack of judgment regarding which programs suit better for each study objective [48]. We expect our study benefits researchers to understand functionality differences between the evaluated AI-based plasmid annotation tools, in the light of artificial intelligence as an emerging, helpful, and fast-developing software programming method.

## IV. Conclusions

Our results expose Platon had the best performance among the analyzed plasmid prediction software based on artificial intelligence. We consider that research in the environmental metagenomic area could greatly benefit from the sensitivity, precision, and accuracy of Platon. Nonetheless, RFPlasmid represents a good less-sensitive alternative for plasmidomes annotation in metagenomes datasets and might be helpful in taxon-dependent prediction, which has been reported to outperform the generic approach here studied. Moreover, ARGs prediction indicated the presence of genes encoding resistance against a wide range of antimicrobial compounds, including some of which have been previously reported to be present in Virilla River through chemical annotation, indicating an ecological and clinical risk. For future research, the performance analysis of AI-based and regular annotation tools might uncover a more dense and precise comparison between bioinformatic workflows. Also, the typification and mobility evaluation of plasmids with bioinformatic tools such as MOBsuite [49] can evaluate the similarity among prediction tools output. Finally, the study of more metagenomic samples of the Virilla River will provide insights into a more trustful plasmidome composition, considering the difference annotated.

## V. Acknowledgment

This research was partially supported by a machine allocation on Kabré supercomputer at the Costa Rica National High Technology Center.

## Footnotes

1 https://www.tensorflow.org/

2 https://klif.uu.nl/rfplasmid/

3 https://card.mcmaster.ca/analyze/rgi

