## Supplementary Material for "Benchmarking AI-based plasmid annotation tools for antibiotic resistance genes mining from metagenome of the Virilla River, Costa Rica"


Dorian Rojas-Villalta
*Advanced Computing Laboratory*
*Costa Rica National High Technology Center (CeNAT)*
San José, Costa Rica


Melany Calderón-Osorno
*Costa Rica National High Technology Center (CeNAT)*
San José, Costa Rica


Kenia Barrantes-Jiménez
*Doctorado en Ciencias Naturales para el Desarrollo (DOCINADE)*
*Instituto Tecnológico de Costa Rica, Universidad Nacional, Universidad Estatal a Distancia*
*Instituto de Investigaciones en Salud*
*Universidad de Costa Rica*
San José, Costa Rica


Maria Arias-Andres
*Instituto Regional de Estudios en Sustancias Tóxicas*
*Universidad Nacional*
Heredia, Costa Rica


Keilor Rojas-Jiménez
*Escuela de Biología*
*Universidad de Costa Rica*
San José, Costa Rica




979-8-3503-3005-2/23/$31.00 ©2023 IEEE

TABLE I

ANTIBIOTIC RESISTANCE GENES (ARGS) PREDICTION OF WHOLE METAGENOME, PLASFLOW, PLATON, RFPLASMID AND PLASFOREST ANNOTATED PLASMIDOMES

| Sample | ARO Term | AMR Gene Family | Drug Class | Resistance mechanism | Identity of Matching Region (%) |
|---|---|---|---|---|---|
| Whole metagenome | *qacG* | Small multidrug resistance antibiotic efflux pump | Disinfecting agents and antiseptics | Antibiotic efflux | 44.55 |
| | *adeF* | Resistance-nodulation-cell division antibiotic efflux pump | Fluoroquinolone and tetracycline antibiotics | Antibiotic efflux | 42.91 |
| | *adeF* | Resistance-nodulation-cell division antibiotic efflux pump | Fluoroquinolone and tetracycline antibiotics | Antibiotic efflux | 45.6 |
| | *adeF* | Resistance-nodulation-cell division antibiotic efflux pump | Fluoroquinolone and tetracycline antibiotics | Antibiotic efflux | 43.31 |
| | *vanT* gene in *vanG* cluster | Glycopeptide resistance gene cluster *vanT* | Glycopeptide antibiotic | Antibiotic target alteration | 43.3 |
| | *adeF* | Resistance-nodulation-cell division antibiotic efflux pump | Fluoroquinolone and tetracycline antibiotics | Antibiotic efflux | 43.3 |
| | *adeF* | Resistance-nodulation-cell division antibiotic efflux pump | Fluoroquinolone and tetracycline antibiotics | Antibiotic efflux | 84.46 |
| | *qacG* | Small multidrug resistance antibiotic efflux pump | Disinfecting agents and antiseptics | Antibiotic efflux | 39.42 |
| | *vanT* gene in *vanG* cluster | Glycopeptide resistance gene cluster *vanT* | Glycopeptide antibiotic | Antibiotic target alteration | 32.81 |
| | *qacG* | Small multidrug resistance antibiotic efflux pump | Disinfecting agents and antiseptics | Antibiotic efflux | 38.1 |
| | *qacG* | Small multidrug resistance antibiotic efflux pump | Disinfecting agents and antiseptics | Antibiotic efflux | 39.42 |
| | *adeF* | Resistance-nodulation-cell division antibiotic efflux pump | Fluoroquinolone and tetracycline antibiotics | Antibiotic efflux | 85.57 |
| | *qacJ* | Small multidrug resistance antibiotic efflux pump | Disinfecting agents and antiseptics | Antibiotic efflux | 38.46 |
| | ***sul1*** | **Sulfonamide resistance sul** | **Sulfonamide antibiotic** | **Antibiotic target alteration** | **99.64** |
| | *adeF* | Resistance-nodulation-cell division antibiotic efflux pump | Fluoroquinolone and tetracycline antibiotics | Antibiotic efflux | 84.54 |
| | ***APH(3")-lh*** | **APH(3")** | **Aminoglycoside antibiotic** | **Antibiotic inactivation** | **99.25** |
| | *vanY* gene in *vanA* cluster | *vanY* glycopeptide resistance gene cluster | Glycopeptide antibiotic | Antibiotic target alteration | 29.5 |
| | *aadA27* | ANT(3") | Aminoglycoside antibiotic | Antibiotic inactivation | 98.46 |
| | ***qacEdelta1*** | **Major facilitator superfamily antibiotic efflux pump** | **Disinfecting agents and antiseptics** | **Antibiotic efflux** | **100** |
| | *vanY* genes in *vanB* cluster | *vanY* glycopeptide resistance gene cluster | Glycopeptide antibiotic | Antibiotic target alteration | 28.89 |
| | *qacG* | Small multidrug resistance antibiotic efflux pump | Disinfecting agents and antiseptics | Antibiotic efflux | 42.31 |
| Plasflow annotated plasmidome | *vanT* gene in *vanG* cluster | Glycopeptide resistance gene cluster *vanT* | Glycopeptide antibiotic | Antibiotic target alteration | 32.81 |
| | *adeF* | Resistance-nodulation-cell division antibiotic efflux pump | Fluoroquinolone and tetracycline antibiotics | Antibiotic efflux | 85.57 |
| | ***qacEdelta1*** | **Major facilitator superfamily antibiotic efflux pump** | **Disinfecting agents and antiseptics** | **Antibiotic efflux** | **100** |
| | ***sul1*** | **Sulfonamide resistance sul** | **Sulfonamide antibiotic** | **Antibiotic target alteration** | **99.64** |
| | ***APH(3")-lh*** | **APH(3")** | **Aminoglycoside antibiotic** | **Antibiotic inactivation** | **99.25** |
| | *aadA27* | ANT(3") | Aminoglycoside antibiotic | Antibiotic inactivation | 98.46 |
| Platon annotated plasmidome | ***qacEdelta1*** | **Major facilitator superfamily antibiotic efflux pump** | **Disinfecting agents and antiseptics** | **Antibiotic efflux** | **100** |
| | ***sul1*** | **Sulfonamide resistance sul** | **Sulfonamide antibiotic** | **Antibiotic target alteration** | **99.64** |
| | ***APH(3")-lh*** | **APH(3")** | **Aminoglycoside antibiotic** | **Antibiotic inactivation** | **99.25** |
| | *aadA27* | ANT(3") | Aminoglycoside antibiotic | Antibiotic inactivation | 98.46 |
| RFPlasmid annotated plasmidome | ***qacEdelta1*** | **Major facilitator superfamily antibiotic efflux pump** | **Disinfecting agents and antiseptics** | **Antibiotic efflux** | **100** |
| | ***sul1*** | **Sulfonamide resistance sul** | **Sulfonamide antibiotic** | **Antibiotic target alteration** | **99.64** |
| | ***APH(3")-lh*** | **APH(3")** | **Aminoglycoside antibiotic** | **Antibiotic inactivation** | **99.25** |
| PlasForest annotated plasmidome | ***qacEdelta1*** | **Major facilitator superfamily antibiotic efflux pump** | **Disinfecting agents and antiseptics** | **Antibiotic efflux** | **100** |
| | ***sul1*** | **Sulfonamide resistance sul** | **Sulfonamide antibiotic** | **Antibiotic target alteration** | **99.64** |
| | *adeF* | Resistance-nodulation-cell division antibiotic efflux pump | Fluoroquinolone and tetracycline antibiotics | Antibiotic efflux | 85.57 |
| | ***APH(3")-lh*** | **APH(3")** | **Aminoglycoside antibiotic** | **Antibiotic inactivation** | **99.25** |

Bold text represents those antibiotic resistance genes present among all samples.
